# How do blind people represent rainbows? Disentangling components of conceptual representations

**DOI:** 10.1101/287318

**Authors:** Ella Striem-Amit, Xiaoying Wang, Yanchao Bi, Alfonso Caramazza

**Author notes:** These authors contributed equally to this work (Ella Striem-Amit, Xiaoying Wang). Corresponding authors: Ella Striem-Amit, Department of Psychology, Harvard University, Cambridge, MA 02138 USA,; Yanchao Bi, State Key Laboratory of Cognitive Neuroscience and Learning & IDG/McGovern Institute for Brain Research, Beijing Normal University, Beijing, 100875 China,.

## Abstract

How do we represent information that has no sensory features? How are abstract concepts like “freedom”, devoid of external perceptible referents, represented in the brain? To address the role of sensory information in the neural representation of concepts, we investigated how people born blind process concepts whose referents are imperceptible to them because of their visual nature (e.g. “rainbow”, or “red”). We find that the left dorsal anterior temporal lobe (ATL) shows preference both to typical abstract concepts (“freedom”) and to concepts whose referents are not sensorially-available to the blind (“rainbow”), as compared to partially sensorially-perceptible referents (e.g. “rain”). Activation pattern similarity in dorsal ATL is related to the sensorial-accessibility ratings of the concepts in the blind. Parts of inferior-lateral aspects of ATL and the temporal pole responded preferentially to abstract concepts devoid of *any* external referents (“freedom”) relative to imperceptible objects, in effect distinguishing between object and non-object concepts. The medial ATL showed a preference for concrete concepts (“cup”), along with a preference for partly perceptible items to the blind (“rain”, as compared with “rainbow”), indicating this region’s role in representing concepts with sensory referents beyond vision. The findings point to a new division of labor among medial, dorsal and lateral aspects of ATL in representing different properties of object and non-object concepts.

## Introduction

How do we represent concepts that extend beyond our perceptual experience, concepts like “freedom” and “justice”, which have no clear external referent? And how do blind people represent concepts such as rainbow, whose referent is perceptible only visually and comprised of colors, which are uniquely visual qualia?

Various studies have addressed the neural correlates of concrete and abstract concepts ^1–4^. Because concrete concepts, like “cup”, have perceptible features, such as shape, size and color, whereas abstract concepts, like “freedom”, lack sensory features, it has been proposed that the latter type of concepts rely more heavily on semantic or verbal information ^5,6^. Therefore, the inspection of how abstract concepts are represented has been considered an important way to understand language and knowledge representation in the brain. Traditionally, this has been tested by comparing brain responses to abstract concepts to those generated by concrete ones (like “cup”). This comparison has revealed large-scale networks of regions associated with abstract concepts involving language areas and concrete concepts involving modality-specific areas ^2,7–10^.

However, there are additional differences between abstract and concrete concepts beyond the existence of external sensory referents. Abstract concepts tend to be learned later in life, be less familiar ^11,12^ and some of them refer to emotional contents ^13,14^, potentially providing an emotional (internal) “sensory” referent. Furthermore, abstract concepts are more ambiguous and their interpretation depends more on context-dependent variation ^15^. Therefore, the difference between classical abstract and concrete concepts in terms of their sensory features is confounded by additional factors. Furthermore, abstract and concrete concepts differ in an additional important dimension, beyond their sensory availability: their mere objecthood. Concrete concepts generally refer to external objects or referents which can be “pointed” to in the world, whereas abstract concepts do not. Nevertheless, these two dimensions are impossible to be teased apart in most circumstances as referents are intrinsically sensible.

How can the effects of sensory availability and experience, as well as that of objecthood, be tested then? Here we take a unique approach to overcome these confounds and inspect the roles of these conceptual dimensions directly, by using a special population that does not have access to sensorially perceptible referents for otherwise concrete object concepts, thereby eliminating the confounds mentioned above. To this aim we studied a group of people born blind as they were presented with concepts that have both object referents and sensory-accessible features (“cup”); concepts that have external referents but are perceivable through vision alone, and are thus sensorially-inaccessible objects to the congenitally blind (“rainbow”); and abstract concepts without referents or sensory features (“freedom”). This gradient of concepts between fully concrete and fully abstract concepts in the blind allows us to separate sensory components and objecthood and study their neural correlates.

## Results

To inspect how abstract information is represented in the brain, we first localized brain preference for classical abstract concepts, chosen carefully as to not arouse strong emotional responses (see Table 3). Similarly to previous reports, abstract concepts (“freedom”, compared to concrete every-day objects that are similarly familiar to the blind; “cup”; see Fig. 1A two right-most columns) evoked significant activation in multiple regions, mainly left-lateralized, in the combined subject group (Fig. 1B; for similar findings in each group separately and the reverse contrast see Fig. S1). These included the inferior frontal lobe, superior temporal sulcus and anterior temporal lobe (ATL), both in the anterior superior temporal plane, as well as below it towards the temporal pole. A more stringent contrast, in which the abstract concepts condition was further required to also elicit significant positive activation (abstract > concrete AND abstract > baseline), limited this network to the left hemisphere, and within the ATL, mainly to the dorsal and lateral aspects (Fig. 1C).

**Figure 1:**
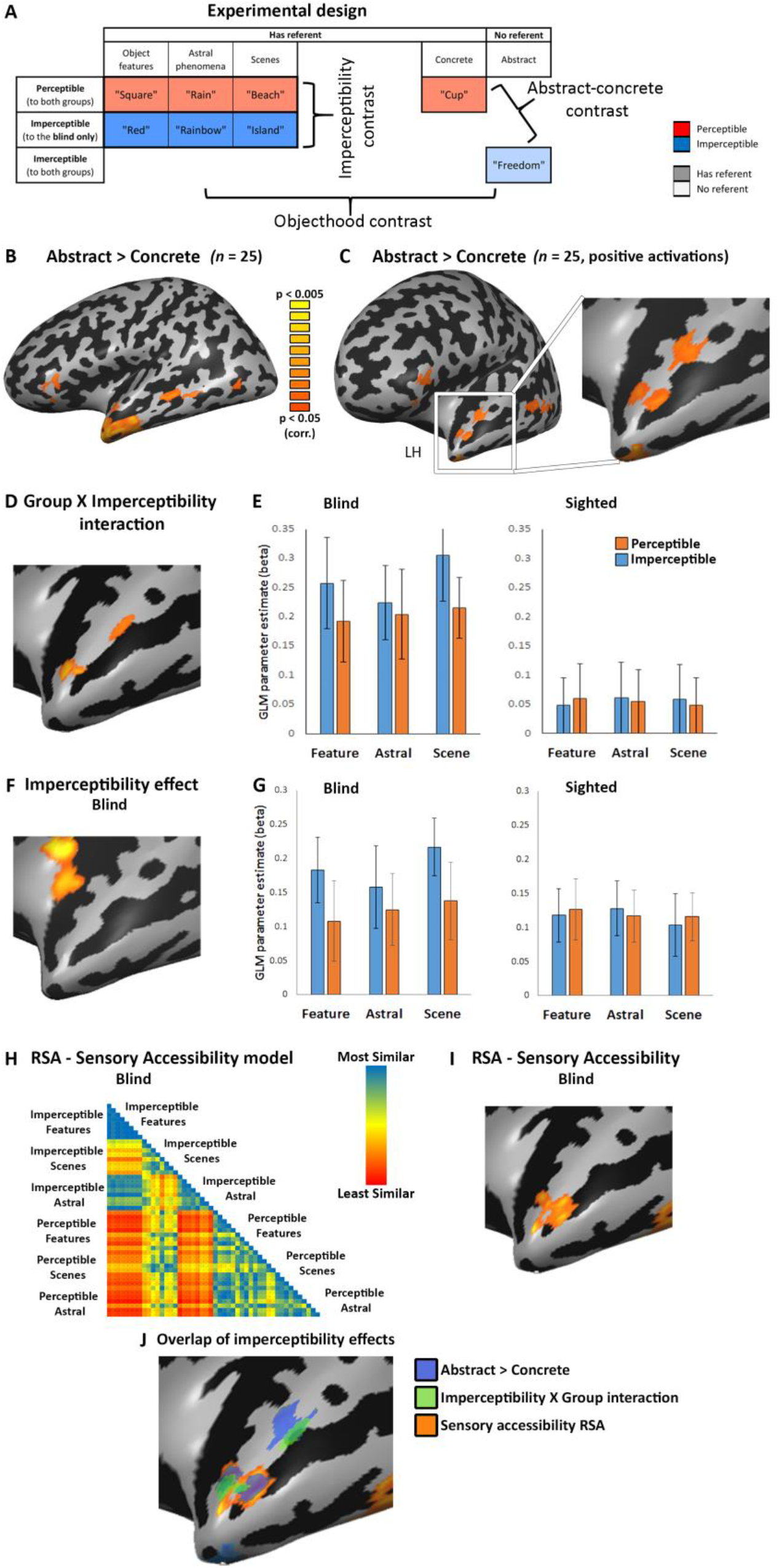
Imperceptible concepts processing is supported by the left dorsal ATL. **A.** The experimental design is depicted, along with examples of the stimuli. The row effect is that of perceptibility: items which are either at least partially perceptible to both blind and sighted (red color), imperceptible completely to the blind (blue color), or Imperceptible to both groups (light blue). The column effect is that of objecthood. The first four columns have external referents (dark colors), whereas the fifth one (abstract concepts; light color) does not. Within the first three columns, different content domains of concepts which have external referents are shown (object features, astral phenomena and scenes). For the imperceptibility contrast, all three content domains are compared between imperceptible and perceptible concepts (dark blue vs. dark red). For the objecthood contrast, imperceptible concepts without referent (abstract concepts, light blue) are compared with imperceptible concepts with referents (dark blue). **B.** A contrast of typical abstract words (e.g. “freedom”) with concrete everyday objects (e.g. “cup”) in the combined subject group (n=25) shows a left-lateralized fronto-parietal network consistent with previous findings ^2,7,9^. **C.** A more stringent contrast requiring the abstract concepts to also generate significantly positive activation focuses the activation to the left hemisphere, and in the ATL to its dorsal and lateral aspects. This contrast is also presented in an anterior view, focusing on the anterior temporal lobe (ATL). **D.** To probe for the effect of sensory feature accessibility, we compared brain activity in people blind from birth and sighted controls, in response to concepts which have external referents in the world, but are perceptible only through the visual sensory modality, and are thus imperceptible to a blind person (e.g. “rainbow”) as compared to concepts whose referents are sensorially perceptible also to the blind (through other modalities; e.g. “rain”). An area which is sensitive to imperceptibility of concepts should respond differently in the two groups for this contrast, as visually-dominant concepts are fully perceptible to the sighted subjects. The ANOVA effect of Group X Imperceptibility interaction across content domains shows two clusters in ATL which respond differently in the blind and sighted to the presented words based on their perceptibility. **E.** The anterior cluster shown in Fig. 1D shows a preference for imperceptible concepts across concept domains (object features, astral phenomena and scenes) only in the blind group. **F.** Imperceptibility effect, differential activation for perceptible vs imperceptible concepts in the blind group (across content domains), affects only the left dorsal ATL among the areas showing preference for abstract concepts. **G.** The dorsal ATL imperceptibility cluster (shown in Fig. 1F) specifically shows a preference for imperceptible concepts across concept domains (object features, astral phenomena and scenes) only in the blind group. This region therefore prefers concepts without tangible sensory properties. **H. H-I**. Multivariate representational similarity analysis (RSA) was computed comparing a behavioral matrix based on ratings of the blind subjects of the sensory perceptibility of the concepts (**H**) with the neural patterns in a searchlight manner across the brain. Sensory accessibility correlation (**I**) was found in the dorsal ATL, overlapping the effects of imperceptibility X group interaction and abstract concepts preference. **I.** The main effects from Fig. 1 are shown together, to reveal the overlap in the dorsal ATL between preference for abstract concepts (over concrete ones; Fig. 1C; depicted in blue), Imperceptibility X Group interaction (Fig. 1D; depicted in green) and the sensory accessibility RSA (Fig. 1I; depicted in orange).

### Impe*rceptibility – Dorsal ATL*

We then investigated which of those regions were indeed sensitive to the absence of sensory information, as opposed to sensitivity to the existence of external referents or to other confounding factors. To do so, we examined brain activity in people blind from birth (Table 1) for concepts that have external referents in the world, but are perceptible only through the visual sensory modality, and are thus imperceptible, devoid of sensory correlates, to a congenitally blind person (e.g. “rainbow”). We compared these concepts to other concepts from the same content domain (in the case of rainbow, astral/weather phenomena) which also have external referents with sensory features in other senses, and are thus sensorially available, perceptible, to the blind (for example, “rain”; sensory accessibility was rated by blind subjects; see methods). Imperceptible and perceptible concepts were chosen from three different content domains to avoid domain-specific effects: astral/weather phenomena (e.g. “rainbow” vs. “rain”), scenes (“island” vs. “beach”) and object features (colors vs. shapes, e.g. “red” vs. “square”). Importantly, the imperceptibility comparison – ANOVA of the full design, comparing the perceptible and imperceptible concepts across domains; see Fig. 1A – comparing dark red and blue across the first three left-most columns – did not significantly differ in any of the various potentially confounding factors: general concreteness/abstractness, imageability, age of acquisition, familiarity, semantic diversity, emotional valence or arousal (F < 3.25, p > 0.08 for all seven comparisons, for stimuli ratings see Table 2, for complete post-hoc t-test results see Table 3).

**Table 1:** Blind subjects characteristics.

| Subject | Gender | Age | Years of Education | Handedness | Cause of blindness | Light perception |
| --- | --- | --- | --- | --- | --- | --- |
| B1 | M | 36 | 12 | Bi | Congenital microphthalmia | None |
| B2 | M | 22 | 15 | R | Congenital microphthalmia | None |
| B3 | M | 33 | 12 | R | Congenital microphthalmia; microcornea | None |
| B4 | M | 48 | 12 | R | Congenital glaucoma | None |
| B5 | F | 46 | 9 | R | Congenital glaucoma | None |
| B6 | M | 40 | 12 | R | Congenital leukoma | Faint |
| B7 | F | 50 | 12 | R | Cataracts; congenital eyeball dysplasia | Faint |
| B8 | M | 57 | 12 | R | Congenital eyeball dysplasia | None |
| B9 | F | 43 | 12 | R | Congenital glaucoma | None |
| B10 | M | 48 | 12 | R | Congenital microphthalmia; cataracts; leukoma | None |
| B11 | M | 63 | 9 | R | Congenital glaucoma; leukoma | None |
| B12 | F | 41 | 12 | R | Congenital optic nerve atrophy | Faint |

**Table 2:** Stimulus ratings (average across all 12 words in each category)

|  | Concreteness/<br>Abstractness | Familiarity | Age of acquisition | Imageability | Emotional arousal | Emotional valence | Semantic Diversity |
| --- | --- | --- | --- | --- | --- | --- | --- |
| Objects | 6.50 | 6.25 | 4.29 | 6.75 | 5.2 | 3.2 | 1.46 |
| Imperceptible astral | 5.75 | 5.75 | 4.38 | 6.08 | 5.4 | 3.6 | 1.52 |
| Perceptible astral | 5.50 | 5.42 | 4.58 | 5.75 | 3.8 | 4.1 | 1.52 |
| Imperceptible scenes | 6.08 | 5.08 | 5.17 | 6.08 | 5.1 | 3.7 | 1.42 |
| Perceptible scenes | 6.00 | 5.50 | 4.63 | 6.00 | 4.8 | 3.6 | 1.62 |
| Imperceptible features (Colors) | 4.75 | 5.42 | 4.63 | 5.83 | 6.2 | 3.4 | 1.71 |
| Perceptible features (Shapes) | 5.50 | 5.83 | 4.63 | 6.08 | 6.4 | 4 | 1.62 |
| Abstract | 1.92 | 5.92 | 4.92 | 2.00 | 3.9 | 3.1 | 1.82 |

**Table 3:** Stimulus differences. (student’s t-test p values)

| Contrast | Concreteness/<br>Abstractness | Familiarity | Age of acquisition | Imageability | Emotional arousal | Emotional valence | Semantic Diversity |
| --- | --- | --- | --- | --- | --- | --- | --- |
| Abstract vs. concrete | 0.000* | 0.22 | 0.11 | 0.000* | 0.06 | 0.84 | 0.000* |
| Overall imperceptible vs. perceptible design<sup>1</sup> | 0.17 | 0.48 | 0.53 | 0.79 | 0.13 | 0.12 | 0.01 |
| Abstract vs. imperceptible astral | 0.000* | 0.67 | 0.30 | 0.000* | 0.03 | 0.27 | 0.03 |

\* Statistically significant; Bonferroni corrected for multiple comparisons.
<sup>1</sup> The overall imperceptible vs. perceptible design includes the comparison between the imperceptible and perceptible words in the three tested domains: astral/weather phenomena (e.g. “rainbow” vs. “rain”), scenes (“island” vs. “beach”) and object features (colors vs. shapes, e.g. “red” vs. “square”)

Given that the imperceptible concepts are only sensorially inaccessible to the blind, we expected regions differentially engaged due to the sensory imperceptibility of concepts to show differential responses for the blind and sighted subjects in our experimental design. We therefore computed an ANOVA model for a domain X imperceptibility X group effect, and looked for areas showing a group X imperceptibility interaction, a different response based on perceptibility in the two groups, across concept domains. Among the brain regions showing preference for abstract concepts in both groups, only the left ATL also showed such an interaction, in two clusters in the superior ATL (Fig. 1D; see the overlap between these contrasts in Fig. 1J). In the anterior cluster of this map, in left dorsal ATL, this interaction manifested in heightened activity in the blind group for imperceptible concepts across the three content domains (Fig. 1E, data sampled in an independent dataset (event-related design; see methods) from anterior STG cluster in Fig. 1D. For detail of the posterior cluster see detail in the next section). A significant effect of imperceptibility in the blind group, in a simplified imperceptibility X domain ANOVA model, was found in a slightly more dorsal part of ATL, in the anterior superior temporal plane (Fig. 1F). Specifically, this superior ATL imperceptibility cluster shows a main effect of imperceptibility in the blind (F = 8.63, p < 0.05; see sampled data in Fig. 1G), a significant preference for imperceptible concepts, but no domain effect (p > 0.4) or interaction (p > 0.61), signifying that Sensory accessibility affects this region beyond the divergent content domains. Importantly, the sighted group showed no such effect in this region (all effects and interaction p > 0.82), and a combined ANOVA with both groups in this region revealed an imperceptibility X group interaction (F = 5.46, p < 0.05), supporting the absence of vision as the factor behind the imperceptible/perceptible category differences.

The univariate analyses reported here show a preference for imperceptible concepts in left dorsal ATL. Converging evidence from multivariate analyses further supports the role of imperceptibility in determining concept property preferences in dorsal ATL. Using behavioral ratings of the perceptible and imperceptible objects in the congenitally blind group, we computed a dissimilarity matrix of the stimuli based on their perceptual accessibility (Fig. 1H). A multivariate comparison of the neural similarity matrices from the single-item-level event-related data in the blind, with this model of imperceptibility (searchlight representational similarity analysis; RSA), shows that the anterior dorsal ATL response pattern indeed varies based on this parameter (Fig. 1I). This cluster overlaps the area showing the abstract > concrete effect as well as the group X imperceptibility interaction (Fig. 1D; see overlap in Fig. 1J). As these ratings do not reflect the sensory accessibility of the sighted, it is not surprising that no similar correlation between this behavioral matrix is found for the sighted neural data. Therefore, evidence from both univariate and multivariate analyses support the role of left dorsal ATL in processing imperceptible concepts in the blind, suggesting that this region’s response to abstract concepts is affected by the absence of sensory information regardless of objecthood (as both perceptible and imperceptible concepts have external referents) and other confounding factors.

### Objecthood – Lateral ATL

The results show that lateral aspects of left ATL also prefer abstract concepts (Fig. 1C). However, the posterior peak showing the imperceptibility X group interaction (Fig. 1D) does not seem to be associated with a significant imperceptibility effect in the blind group, in either the univariate or multivariate analyses (Fig. 1F-I). To clarify this region’s role in abstract concept processing, we further investigated its responses to perceptible and imperceptible concepts in the blind. Sampling the inferior lateral imperceptibility X group interaction cluster, we find that this region does not show a clear imperceptibility effect or preference in the blind, but rather inconsistent responses for different conceptual domains (Fig. 2A; the imperceptibility X group interaction arises from a bias in the sighted group; Fig. S2).

**Figure 2:**
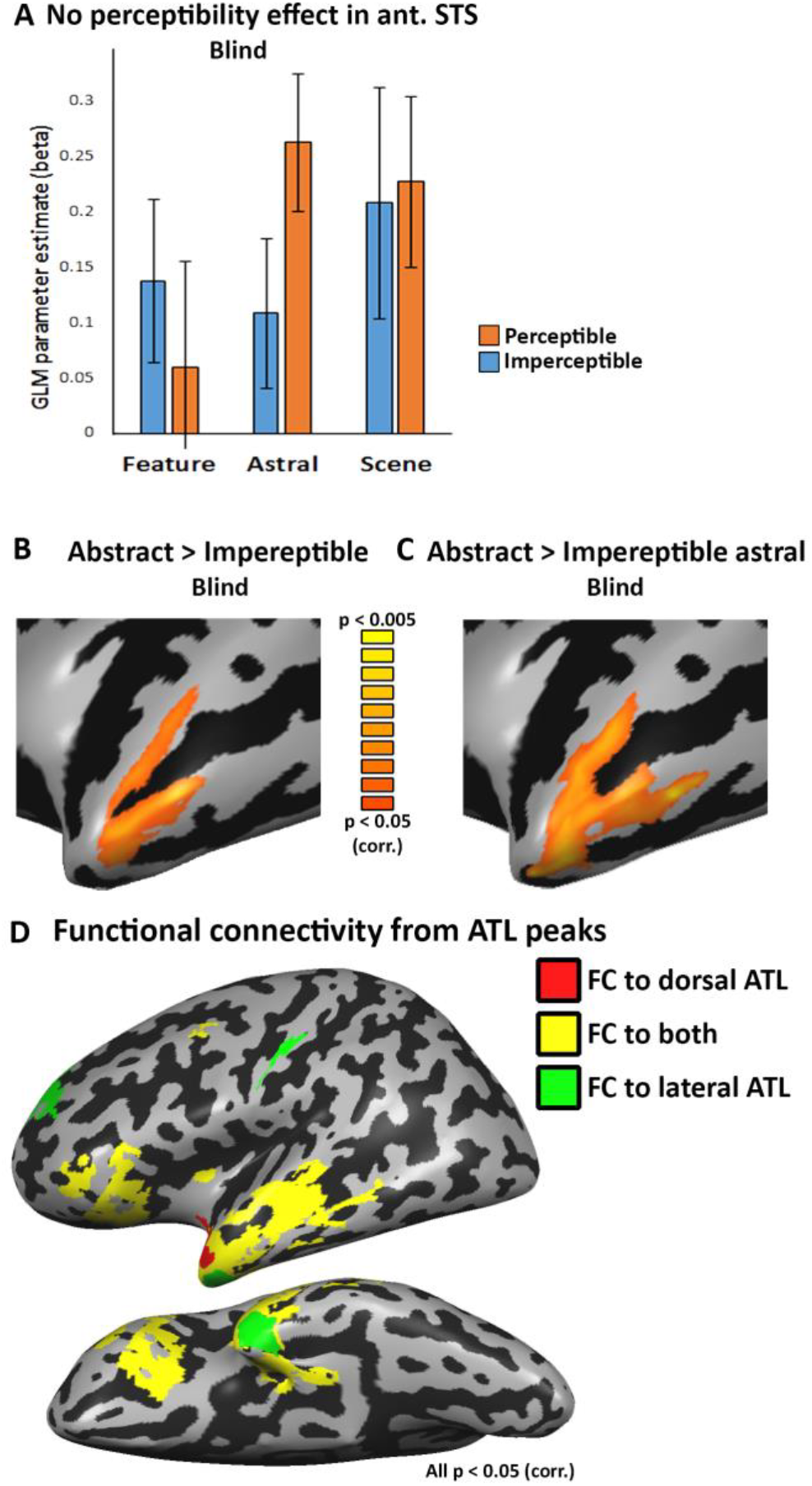
Lateral ATL shows preference for concepts without external referents. **A.** The ANOVA effect of Group X imperceptibility interaction (Fig. 1D) showed also a lateral ATL cluster (in anterior STS, the posterior cluster in Fig. 1D). However, this region’s responses shown here, sampled in the blind group, show no clear perceptibility effect or a consistent pattern of preference for either perceptible or imperceptible concepts across domains. The interaction with group appears to originate from a bias in the sighted group which does not correlate to perceptibility (Fig. S2). **B.** The lateral ATL shows a preference for abstract, referent-free, concepts (“freedom”) over imperceptible concepts (whose external referents are not sensorially accessible; “rainbow”, “red” and “island”) in the blind, suggesting this region’s preference for abstract concepts relates to the absence of objecthood. No areas showed significant activation for the opposite contrast, preference for imperceptible concepts over abstract ones. **C.** The preference for referent-free concepts in lateral ATL is replicated as compared to the astral imperceptible concepts domain alone (“rainbow”), which are more typical (figurative) objects. **D.** Partial functional connectivity was computed from the dorsal (red) and lateral (green) ATL peaks in the sighted. Overlapping FC to both seeds (in yellow) is predominant, showing that these two regions belong largely the same functional network.

Can this region’s preference for abstract concepts over concrete ones be explained by the other dimension of abstractness, the absence of objecthood, of not having an external referent? To study this possibility, we compared abstract concepts that are both physical referent-free and devoid of sensory correlates (e.g., “freedom”) to concepts that have imperceptible referents in the blind (e.g., “rainbow”; see Fig. 1A; comparing light and dark blue), thus removing the bias of sensory information. This contrast activated the more lateral ATL regions involved in abstract concepts over concrete ones, and extending anteriorly towards the temporal pole (Fig. 2B). As some of our imperceptible concept domains are object features (“red”) and scenes (“island”) rather than classical objects themselves, we additionally replicated this finding using the astral/weather imperceptible concepts such as “rainbow” and “moon”, which are more classical figurative objects ^16^ (Fig. 2C; these concepts are also comparable to abstract concepts in all relevant behavioral measures; see details in methods and Table 3). Therefore, the lateral and anterior ATL’s preference for abstract concepts over concrete ones (“freedom” over “cup”; in Fig. 1B) seems to result from a preference for referent-free concepts, even within imperceptible concepts. Interestingly, the effect of objecthood overlapped to some extent with areas showing the effect of imperceptibility and its interaction between the groups, suggesting that these two dimensions are not completely orthogonal. Instead, the overlap area on the upper banks of the anterior superior temporal sulcus appear to be affected by both factors.

We also inspected if the two regions would belong to the same or different functional networks. We computed functional connectivity from seeds at the peaks of the cluster showing the group X imperceptibility interaction in the dorsal ATL (Fig. 1D) and the peak of the cluster showing the abstract > imperceptible concepts in the ATL pole (Fig. 2B). Despite their difference in functional preferences, the dorsal and lateral ATL seem to belong largely to the same functional network, which includes large parts of the dorsolateral ATL and inferior frontal lobe (Fig. 2D; note the prevalence of shared FC marked in yellow). The spatial overlap and shared network suggest that these regions may be part of the same system for the processing of semantic, non-sensorially derived information.

### “Concrete” concepts – Medial ATL

What of the reverse effect, of having sensorially accessible properties? The contrast of concrete objects (“cup”) vs. abstract concepts (“freedom”) highlighted a known network of multisensory object processing ^17–19^, including the medial ATL (perirhinal cortex) in both groups (Fig. 3A; see also separately in the sighted group; Fig. 3B; Fig. S1). If this region has a role in processing sensory features of objects, we can expect to find a preference for perceptible (“rain”) over imperceptible (“rainbow”) concepts in the blind, and this is indeed the case in the medial ATL (Fig. 3C; across all three content domains; comparing dark blue and red across the first three left-most columns in Fig. 1A). Therefore, the medial ATL appears to have an opposite preference than the dorsal ATL. We further investigated if this dissociation of preferences in these two regions would also manifest in having different network connectivity patterns by plotting their partial functional connectivity (FC; Fig. 3D). Seeds were selected from the peaks of the cluster showing the group X imperceptibility interaction in the dorsal ATL (shown in Fig. 1D; also used for Fig. 2D) and of the cluster showing the preference for concrete vs. abstract concepts in medial ATL (shown in Fig. 3A). The partial FC shows that the medial ATL is better connected to multisensory object-related regions in the frontal lobe, parietal lobe, as well as in the ventral visual cortex. This connectivity profile is consistent with the literature linking medial structures in ATL, mainly the perirhinal cortex, as the mechanism of sensory feature integration of object features ^20–22^. In contrast, the dorsal ATL is more strongly connected to the lateral and anterior ATL towards the temporal pole, as well as to the inferior frontal lobe, parts of the language network. Therefore, the FC analysis also supports the distinct roles of these subparts of ATL.

**Figure 3:**
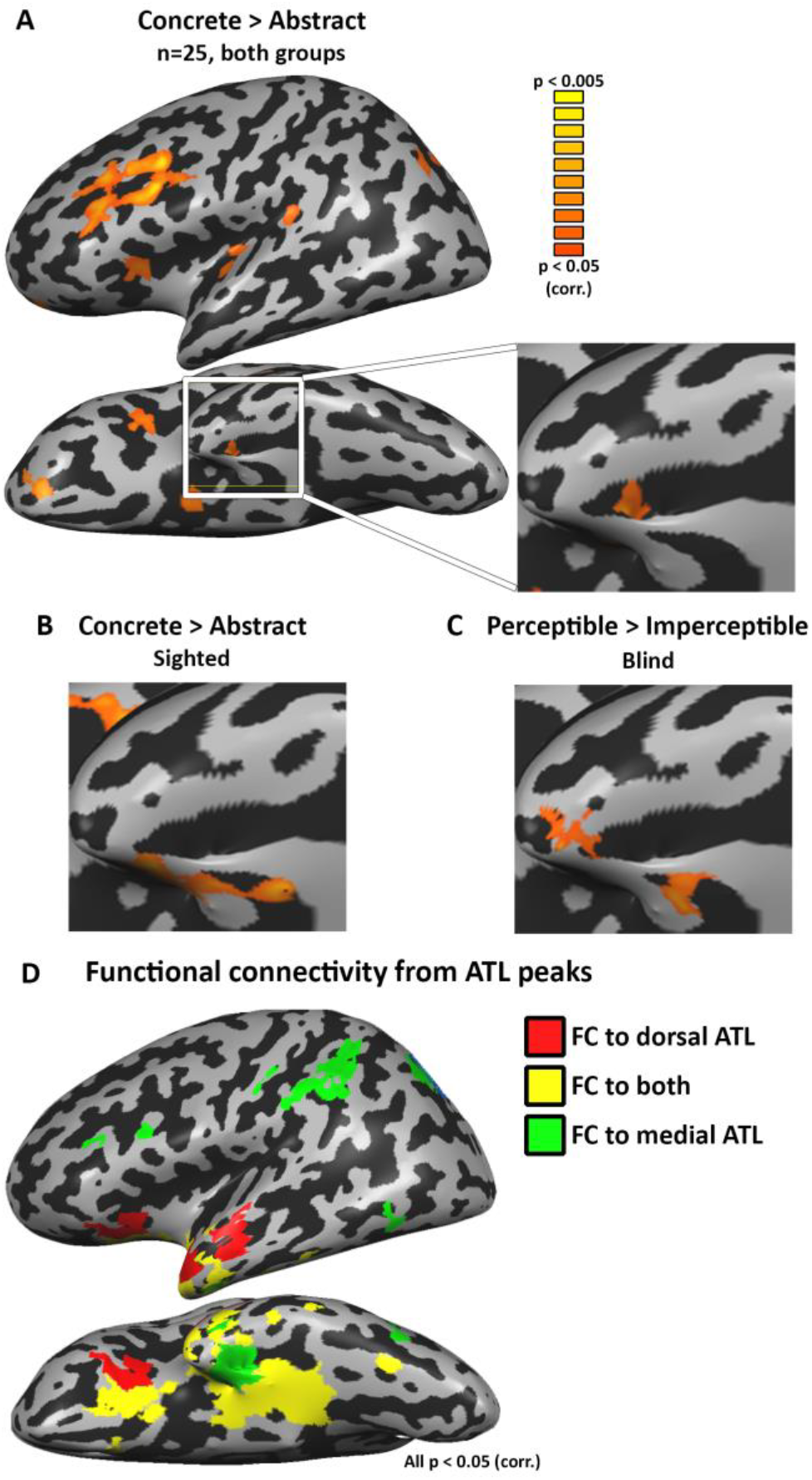
Perceptible concepts processing is supported by the medial ATL. **A-B.** A contrast of concrete everyday objects as compared with typical abstract words in the combined subject group (n=25, in **A**; see **B** for sighted group separately) shows a network of regions associated with multisensory object perception. In the ATL, the medial ATL shows preference for processing concrete objects. **C.** The medial ATL also shows a clear preference for perceptible (e.g. “rain”) over imperceptible (“rainbow”) concepts in the blind, although the concepts are perceptible via non-visual modalities. **D.** The dorsal and medial ATL regions which show opposite preferences for perceptibility in the blind belong to largely different functional networks in the sighted. Partial functional connectivity is plotted for the dorsal (red) and medial (green) ATL. Overlapping FC to both seeds is depicted in yellow.

## Discussion

We find that the response of various parts of the ATL to abstract concepts can be broken down into effects of imperceptibility and of objecthood. Words devoid of sensorially-accessible, tangible features, either classical abstract concepts (“freedom”) or words depicting visually-dominant phenomena (“rainbow”) in blind people, show preferred activation in the left dorsal superior ATL (Fig. 1). Supporting evidence for this are the results of the multivariate representational similarity analysis (RSA) which found that the activation pattern in this region correlated negatively with the level of sensory accessibility of the concepts in the blind (Fig. 1H). In contrast, the lateral areas in anterior STS and the temporal pole show a preference for abstract concepts without a corresponding preference for imperceptible concepts in the blind (Fig. 2A). Instead, their activation in response to abstract concepts exceeds that of even imperceptible concepts, suggesting a role for the absence of referents altogether – the absence of objecthood – in determining this regions’ representational preference (Fig. 2B, C). These two regions are strongly functionally connected (Fig. 3D), suggesting parallel involvement in processing different but similarly amodal contents of conceptual information. Our results also further support the role of the medial ATL, the perirhinal cortex and nearby regions, in processing sensorially derived properties of concepts, as it shows a combined preference for concrete objects as well as perceptible objects in the blind (Fig. 3A, B, C). This medial aspect of ATL is more strongly connected functionally to multisensory object-processing regions than the dorsal and lateral aspects of ATL (Fig. 3D). The findings reported here reveal a richly articulated neural organization of the various dimensions of the representation of abstract and concrete concepts.

First, these findings support the role of ATL in processing semantic content related to aspects of sensorially-derived properties, including objecthood, while controlling for common confounds associated with the typical items used to evaluate the representation of abstract versus concrete concepts ^2,11–13^. Multiple neuroimaging studies have emphasized the role of the superior, dorsolateral ATL, and specifically the anterior STG, in the representation and retrieval of semantic and conceptual information ^1,4,7,10,23–26^. Furthermore, studies of the temporal variant of frontotemporal dementia, which results in the selective deterioration of semantic knowledge, have implicated the bilateral ATL in encoding semantic knowledge ^1,23,25,27–30^. Our findings support the ATL’s role in the representation of conceptual knowledge and show that the content processed in these regions can be such that it extends beyond sensory experience and referent.

Second, this study reveals functional dissociations within the ATL – dorsal, lateral, and medial aspects of ATL – based on the effects of perceptibility and objecthood. This division is more fine-grained in nature within the dorsolateral cortex. Both dorsal and lateral (middle) ATL show a preference for abstract over concrete concepts, linking them to abstract conceptual knowledge. The ATL’s most dorsal aspect showed a preference for abstract concepts due to their sensory inaccessibility whereas the lateral aspects were sensitive to the absence of object referents altogether. A partial overlap between these two concept types was found in the dorsal banks of the STS, suggesting that the crucial factor is the absence of different aspects of sensory reference. Consistent with this view is the findings of functional connectivity (Fig. 2D), showing that dorsal and lateral ATL are found to belong largely to the same functional network (see also ^31,32^. Therefore, it appears that both regions process abstract conceptual information, one concerning primarily imperceptible *object* concepts, and the other concerning concept domains which do not correspond to objects (but see below for action concepts).

The distinction reported here between the roles of dorsal and lateral ATL in conceptual processing is subtle, reflecting the contribution of different aspects of abstract concepts. Much more substantial are the distinct roles of dorsolateral and ventromedial ATL in conceptual processing, reflecting the distinction between “abstract” and “concrete” concepts, respectively. This finding is in accord with research linking the medial aspect of ATL, and particularly the perirhinal cortex, to processing of sensorially-derived conceptual properties of objects ^1,20,33–36^. The functional dissociation we find between dorsolateral and ventromedial ATL is also in agreement with a the neuropsychological literature dealing with specific deficits for processing concrete concepts more than abstract ones in semantic dementia and some stroke patients – the reverse concreteness effect ^37–39^ and in patients with ATL resection ^40^. Based on our results, such a phenomenon would occur in cases where temporal lobe damage involves the ventro-medial aspect of ATL, sparing (at least initially) its left dorsal aspects. Evidence for medial ATL damage being associated with a deficit in processing sensorially-derived conceptual properties has been demonstrated in semantic dementia patients ^21^.

Our results additionally provide evidence for the role of medial ATL in processing sensorially-derived features of objects beyond vision and visual experience, as they revealed a preference in the blind for processing (non-visually) perceptible objects as opposed to imperceptible ones (Fig. 3). Although, this region’s role has been linked especially to vision and visual representations ^1,41–43^, we found that perceptibility, beyond the visual modality, is the critical component in activating this region. This region is distinct from the lateral and dorsal aspects of ATL and belongs to different functional networks, linking it more robustly to multisensory, object-related regions (Fig. 3D; see also ^32^).

Furthermore, although not tested in our design, there is much evidence that object domain (e.g., animate versus inanimate ^22,44–50^) and other concept properties such as their emotional/social value ^13,51–55^ play a role in the organization of conceptual processing in ATL and more posterior regions of the temporal lobe. In the latter case, predicate-type concepts such as jump, plan, know, and admire involve posterior middle and superior aspects of the temporal lobe ^7,56–61^. Thus, multiple factors contribute to shaping the organization of conceptual information in ATL and the temporal lobe more generally.

To summarize, the approach of using a sensorially-deprived population (the blind) has allowed us to disentangle major components of object and property conceptual knowledge: those related to perceptual properties and representations and those related to non-sensory, modality-independent information. These findings provide evidence for the neural correlates of semantic representations devoid of sensorially-derived features, when controlling for multiple potential confounds, including emotional correlates. This is found across specific content domains in the blind, through both univariate and multivariate analyses, and using both dimensions of sensory accessibility and objecthood. This amodal, sensory-independent level of concept knowledge representation is supported by the dorsolateral ATL. An additional, finer distinction reflects objecthood (e.g., “freedom” versus “rainbow” in the blind) within the larger area representing imperceptible concepts. In contrast, a preference for concrete concepts due to their sensory feature availability is supported by the medial ATL. Thus, the current findings provide important support to the neural dissociation between abstract semantic knowledge and its sensory properties.

## Materials and methods

### Participants

12 congenitally blind and 14 sighted subjects participated in the experiment. Participants in the blind group were between the age of 22 and 63 (mean age = 44.2 years, 8 males), and did not differ from the sighted participants in age or years of education (age: p < 0.85, years of education; p < 0.83). All sighted participants had normal or corrected-to-normal vision. Subjects had no history of neurological disorder. See Table 1 for detailed characteristics of the blind participants. All experimental protocols were approved by institutional review board of Department of Psychology Peking University, China, as well as by the institutional review board of Harvard University, in accordance with the Declaration of Helsinki, and all subjects gave written informed consent.

### Functional Imaging

Images were acquired using a Siemens Prisma 3-T scanner at the Imaging Center for MRI Research, Peking University. The participants lay supine with their heads snugly fixed with foam pads to minimize head movement. Functional imaging data for the main experiment were comprised of four functional runs, each containing 251 continuous whole-brain functional volumes that were acquired with a simultaneous multi-slice (SMS) sequence supplied by Siemens: slice planes scanned along the rectal gyrus, 64 slices, phase encoding direction from posterior to anterior; 2 mm thickness; 0.2mm gap; multi-band factor = 2; TR = 2000 ms; TE = 30 ms; FA = 90°; matrix size = 112 × 112; FOV = 224 × 224 mm; voxel size = 2 × 2 × 2 mm. Functional imaging data for the single-item-level event-related experiment were comprised of eight functional runs, each containing 209 continuous whole-brain functional volumes using the same sequence parameters as the block-design scans. T1-weighted anatomical images were acquired using a 3D MPRAGE sequence: 192 sagital slices; 1mm thickness; TR = 2530 ms; TE = 2.98 ms; inversion time = 1100 ms; FA = 7°; FOV = 256 × 224 mm; voxel size = 0.5 × 0.5 × 1 mm, interpolated; matrix size =512 × 448.

### Experimental paradigm and stimuli

The stimuli for the experiment were spoken words, each a two-symbol word in Mandarin Chinese, belonging to eight concept categories (see Fig. 1A): abstract concepts (e.g. “freedom”), concrete everyday objects (e.g. “cup”), and three additional content domains, astral/weather phenomena, scenes and object features (shape and color names). Those three domains had two different categories each, one which is perceptible through non-visual senses (e.g. “rain”, “beach” and “square”, respectively) and the other which is perceptible only visually (e.g. “rainbow”, “island” and “red”, respectively), and therefore imperceptible to the blind. We used three different content domains for testing the effect of perceptibility such that domain-specific effects would be negligible. Broadly, the visually-dominant categories are those that fit the definition of “figurative”, following the distinction between operative and figurative objects ^16^. Operative objects, used for the perceivable categories here, are defined as those which were relatively discrete and separate from the surrounding context, and easily available to several sense modalities. Figurative elements, in contrast, are those which did not meet these criteria but were nonetheless picturable, and known primarily by their visual configuration. For scenes, operative, perceptible scenes were chosen such that their defining characteristics can be explored non-visually (e.g. “beach”) and figurative, imperceptible, ones chosen to be too large for their overall configuration or defining features to be perceived in non-visual sensory modalities (e.g. “island”). Perceptibility ratings (the extent to which the words have associated sensory information) of the stimuli were collected prior to the experiment by an independent group of six blind subjects who could not participate in the fMRI study. Additionally, the perceptible and imperceptible concepts were rated by the blind fMRI subjects for their sensory accessibility several months after the scan, and were confirmed to be significantly different for all three categories (t-test, p < 0.005 in all three cases).

Each category included 12 words, matched as best as possible for imageability, age of acquisition (AoA), familiarity and concreteness/abstractness, as assessed in an independent sample of 45 sighted Chinese subjects with similar levels of education (see average stimulus ratings in Table 2). Subjects were introduced with each word separately and asked to rate it, in a scale of 1-7 for these characteristics, as reported previously ^62^. Age of acquisition and familiarity are expected to be similar in the sighted and blind subjects for these concepts; even for color concepts which are a uniquely visual qualia, blind adults have shown extensive familiarity ^63,64^, such that they can create an approximated Newton color wheel ^65^, know the colors of everyday objects ^66,67^ (in line with a generally intact vocabulary acquisition;^68^) and only a sensitive similarity measure of one specific concept category (fruit and vegetables) based on color proved to be affected by blindness ^66^. Emotional valence and arousal levels were assessed in a similar manner ^69^ in Mandarin Chinese directly in the blind subjects, several months after the scan. Semantic diversity values were derived from previous literature ^15^. Concrete objects and abstract concepts differed significantly in concreteness/ abstractness, imageability and semantic diversity but not in AoA, familiarity, emotional arousal or valence (see detail for all statistical tests in Table 3). No figurative-operative condition pairs showed significant difference in these parameters (corrected for multiple comparisons; see Table 3). Importantly, the overall imperceptible vs. perceptible design (relevant for fMRI effect depicted in Fig. 1D, F) did not significantly differ in any of the parameters (ANOVA, F < 3.25, p > 0.08). The comparison of the imperceptible astral concepts to the abstract concepts (relevant for fMRI effect depicted in Fig. 2C) differed in the abstractness/concreteness and imageability ratings but not in any other parameter (see Table 3). During the main experiment, the participants kept their eyes closed and heard short lists of words in a block design paradigm (8 second blocks with 8 words each, baseline between blocks 8 seconds). They were instructed to detect and respond to semantic catch trials, a fruit name appearing within blocks (which occurred three times in each run; these blocks were removed from further analysis). Each run began with a 12 sec rest period. Each block contained words from one of the eight concept categories. An item-level slow event-related design was carried out to conduct representational similarity analysis (RSA; ^70^), as well as to be used as an independent dataset for sampling ROI data (as the ROIs were defined from maps plotted from the block-design experiment). The stimuli were eight of the twelve words the perceptible, imperceptible and abstract categories from the main, block-design experiment stimuli. During each of the eight slow event-related runs, the subjects heard each word once, in a random order, followed by a 5 second baseline period. The subjects task was, as in the block-design experiment, to detect fruit names.

### Data analysis

Data analysis was performed using the Brain Voyager QX 2.8 software package (Brain Innovation, Maastricht, Netherlands) using standard preprocessing procedures. The first two images of each scan were excluded from the analysis because of non-steady state magnetization. Functional MRI data preprocessing included head motion correction, slice scan time correction and high-pass filtering (cutoff frequency: 3 cycles/scan) using temporal smoothing in the frequency domain to remove drifts and to improve the signal to noise ratio. No data included in the study showed translational motion exceeding 2 mm in any given axis, or had spike-like motion of more than 1 mm in any direction. Functional and anatomical datasets for each subject were aligned and fit to standardized Talairach space ^71^. Single subject data were spatially smoothed with a three-dimensional 6 mm full-width at half-maximum Gaussian in order to reduce inter-subject anatomical variability, and then grouped using a general linear model (GLM) in a hierarchical random effects analysis (RFX; ^72^). Group analyses were conducted for the blind and sighted group separately (e.g. Fig. S1) and for the combined blind and sighted subject group (n=25, e.g. Fig. 1B, C). Apart from the first comparison of abstract vs. concrete concepts (Fig. 1B), all GLM contrasts between two conditions included comparison of the first term of the subtraction to baseline (rest times between the epochs), to verify that only positive BOLD changes would be included in the analysis. An ANOVA model was computed for group, stimulus domain and perceptibility, including the perceptible and imperceptible stimuli for the object features, scenes and weather/astral phenomena (Fig. 1D; comparing dark red and dark blue three left-most columns in Fig. 1A). Simplified imperceptibility X domain models were computed for the blind group separately (Fig. 1F). The minimum significance level of all results presented in this study was set to p<0.05 corrected for multiple comparisons, using the spatial extent method based on the theory of Gaussian random fields (GRF; ^73,74^ a set-level statistical inference correction). This was done based on the Monte Carlo stimulation approach, extended to 3D datasets using the threshold size plug-in for BrainVoyager QX. The correction was applied in the entire cortex for the abstract vs. concrete (and vice versa; e.g. Fig. 1A, 3A), and for the anatomically defined left ATL (the temporal lobe anterior to Heschl’s gyrus) for the rest of the analyses which focused on this region. To assess the different conditions contribution to the imperceptibility effect, we also sampled the activation GLM parameter estimates for each group and experimental condition in the region (Fig.1D, F) from the independent event-related experiment, showing the imperceptibility effect in the blind in a region-of-interest (ROI) group level random effect analysis in both groups (Fig. 1E, G).

RSA ^70^ from the event-related data was computed as using CoSMoMVPA, an toolbox in Matlab ^75^. Dissimilarity matrices were built from behavioral ratings of the stimuli for their sensory accessibility, as rated by the blind subjects several months after the scan (Fig. 1H). Searchlight pattern correlation analysis was computed for the unsmoothed neural patterns in the blind (from each subjects’ individual behavioral data) and sighted controls (from the median ratings of the blind group). The mean Fisher-transformed correlation for each participant was entered into a one-tailed one-sample t-test against the correlation expected by chance (0) for each group. The resulting map (Fig. 1I) was corrected for multiple comparisons using the spatial extent method, as described above.

### Functional connectivity data analysis and MRI acquisition

A dataset of spontaneous BOLD fluctuations for the investigation of intrinsic (rest state; ^76^) functional connectivity was collected while the subjects lay supine in the scanner without any external stimulation or task. Data was comprised of one functional run, containing 240 continuous whole-brain functional volumes that were acquired with the same EPI sequence and parameters as the main experiment. The first two images of each scan were excluded from the analysis because of non-steady state magnetization. After registration to individual anatomies in Talairach space, ventricles and white matter signal were sampled using a grow-region function embedded in the Brain Voyager from a seed in each individual brain. Using MATLAB (MathWorks, Natick, MA) ventricle and white matter time-courses were regressed out of the data and the resulting time course was filtered to the frequency band-width of 0.1-0.01 Hz (in which typical spontaneous BOLD fluctuations occur). The resulting data were then imported back onto Brain Voyager for further analyses. Single subject data were spatially smoothed with a three-dimensional 6 mm half-width Gaussian. Seed regions-of-interest (ROIs) were defined from the group-level analyses of the task-data. Functional connectivity was computed from (1) a cluster showing group X imperceptibility interaction in the dorsal ATL (Fig. 1D), (2) a cluster showing preference for concrete vs abstract concepts (“cup” vs. “freedom”) in medial ATL (Fig. 3A), (3) a cluster in lateral anterior ATL showing preference to concepts without external referents (“freedom”) over imperceptible astral ones (“rainbow”; Fig. 2C). Individual time courses from these seed ROIs were sampled from each of the sighted participants, z-normalized and used as individual predictors in group random-effect GLM analysis. Partial correlation was also computed for seeds 1 and 2 (Fig. 3D), and seeds 1 and 3 (Fig. 2D), to observe the common and separate networks to which these seeds belong.

## Acknowledgments

We are thankful to the blind subjects who participated in our experiment. This work was supported by National Natural Science Foundation of China (31500882 to X.Y.W., 31671128 to Y.B.); Società Scienze Mente Cervello–Fondazione Cassa di Risparmio di Trento e Rovereto, by a grant from the Provincia Autonoma di Trento, and by a Harvard Provostial postdoctoral fund (to A.C.); and by the European Union’s Horizon 2020 Research and Innovation Programme under Marie Sklodowska-Curie Grant Agreement 654837 and the Israel National Postdoctoral Award Program for Advancing Women in Science (to E.S.-A.); National Program for Special Support of Top-notch Young Professionals (to Y.B.); the Fundamental Research Funds for the Central Universities (2017XTCX04, to Y.B.) and Interdisciplinary Research Funds of Beijing Normal University (to Y.B.).

## Author contribution

E.S.-A., X.W., Y.B. and A.C. designed research; X.W. performed research; E.S.-A. analyzed data; and E.S.-A., X.W., Y.B. and A.C. wrote the paper.

## Supplementary material

**Fig. S1:**
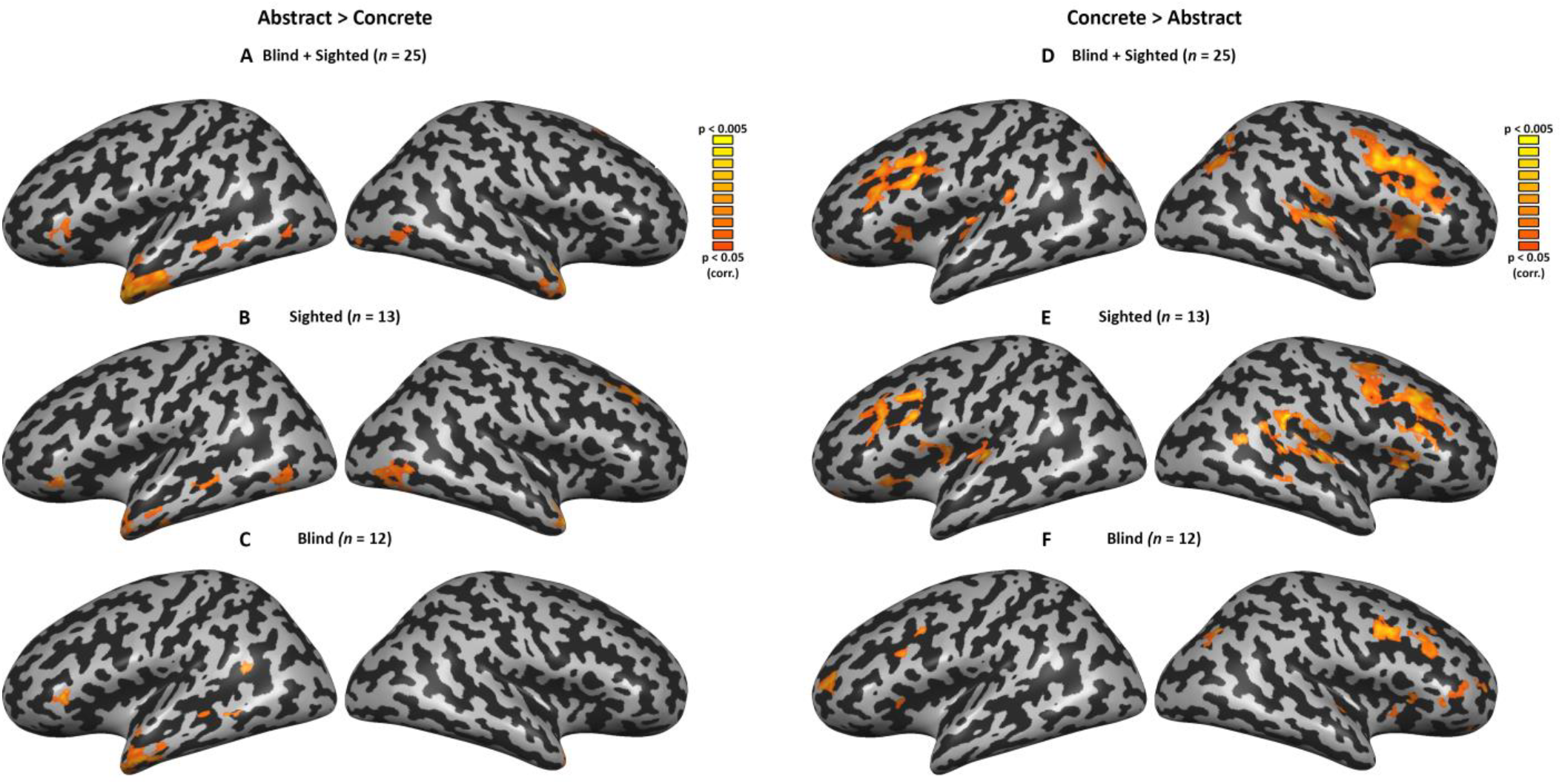
Main effects for abstract concepts and imperceptible concepts in each group. The whole-brain, random effect contrasts of abstract > concrete concepts (**left panels**), concrete > abstract (**right panel**) are depicted for the entire group of subjects (**A,D**) sighted subjects only (**B, E**) and blind subject group (**C, F**).

**Fig. S2:**
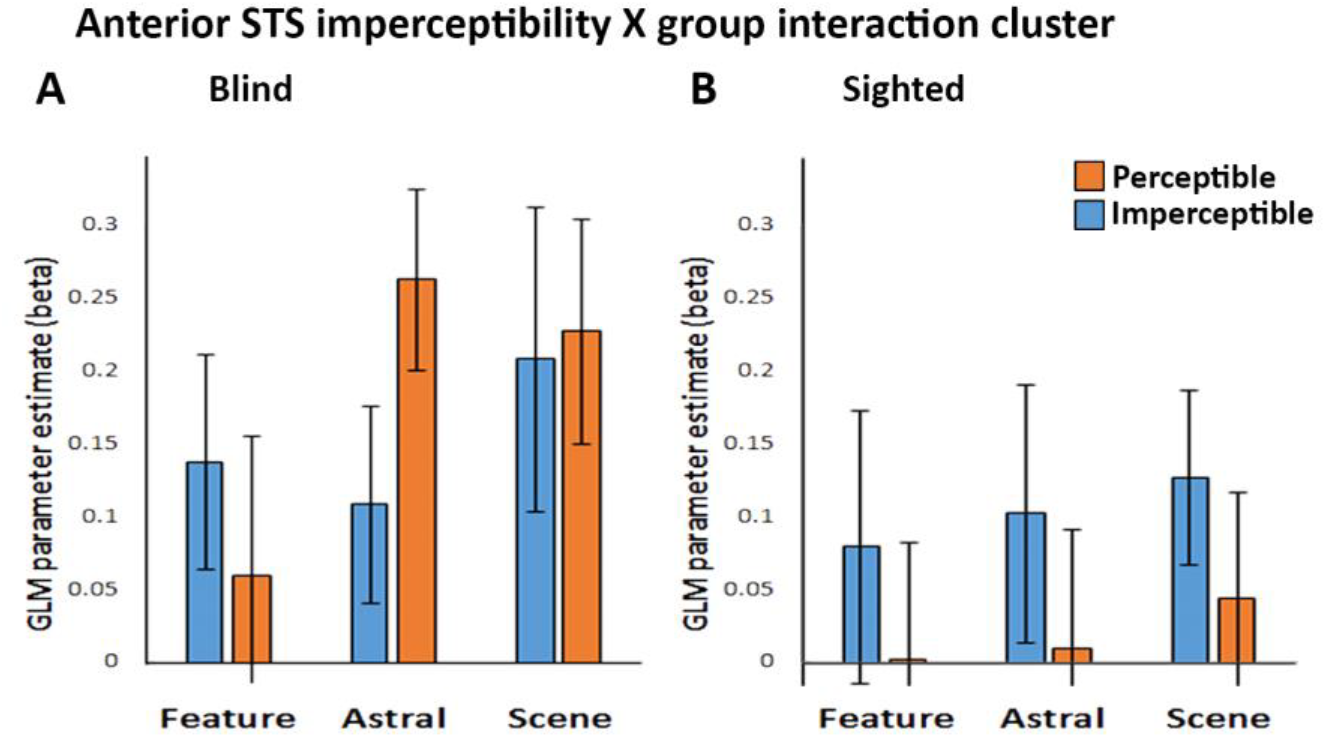
Anterior STS activation pattern. A. The anterior STS imperceptibility X group interaction cluster shown in Fig. 1D does not show a consistent imperceptibility effect across content domains in the blind group. B. The interaction with group appears to result from a preference in the sighted group, towards concepts which are imperceptible to the blind. This cannot be explained in terms of perceptibility, as all these concepts are similarly perceptible to the sighted.

